# Probiotic *Lactiplantibacillus plantarum* VB165 improves metabolic disorders in Insulin-Resistant Mice

**DOI:** 10.64898/2026.03.29.715178

**Authors:** Ting Xu, Wei Zhang, Kexin Jiang, Ting Duan, Xiaoqian Wu, Zirui Zheng, Yanchun Yang, Zhimei Du, Hongmei Zhou, Yan Hui, Shufeng Han, Danqing Chen, Jun Yang

## Abstract

This study investigated the effects of *Lactiplantibacillus plantarum* VB165, a probiotic strain with intrinsic α-glucosidase inhibitor (AGI) activity, on metabolic disorders in high-fat diet (HFD)-induced insulin-resistant (IR) mice. Male C57BL/6 mice were divided into four groups: normal control diet (NCD), NCD supplemented with VB165, HFD, and HFD supplemented with VB165. After 16 weeks, VB165 supplementation significantly attenuated HFD-induced weight gain and reduced epididymal and inguinal white adipose tissue indices. VB165 also improved glucose intolerance and insulin resistance (IR), as demonstrated by oral glucose tolerance tests (OGTT) and insulin tolerance tests (ITT), and lowered fasting blood glucose, fasting insulin, and Homeostatic Model Assessment for Insulin Resistance (HOMA-IR) levels. Additionally, it ameliorated dyslipidemia by reducing serum total cholesterol, triglycerides, and low-density lipoprotein cholesterol (LDL-C), while alleviating hepatic steatosis and adipocyte hypertrophy. Mechanistically, VB165 enhanced intestinal barrier function by upregulating tight junction proteins (ZO-1 and Occludin), reduced systemic inflammation by lowering LPS, IL-6, and IL-1β. Gut microbiota analysis revealed that VB165 modulated community composition, suppressing HFD-enriched genera (e.g., *Ileibacterium* and *Coriobacteriaceae*_UCG_002) and promoting beneficial taxa (e.g., *Faecalibaculum* and *Oscillibacter*). These findings demonstrate that *L. plantarum* VB165 improves HFD-induced metabolic disorders via multi-target mechanisms, highlighting its potential as a probiotic intervention for IR and related metabolic diseases.

## Introduction

Diabetes mellitus represents a major global public health challenge, characterized by chronic hyperglycemia resulting from defects in insulin secretion, insulin action, or both. According to the International Diabetes Federation (IDF) Diabetes Atlas (11th edition, 2025), approximately 589 million adults worldwide were living with diabetes in 2024, a number projected to rise significantly by 2050 ^1^. In China, the diabetes epidemic is particularly severe. Recent estimates indicate that approximately 233 million people were living with diabetes in 2023, representing a prevalence of 15.88% in the total population ^2^. Major drivers include rapid urbanization, dietary transitions towards high-calorie and processed foods, increased sedentary behavior, and rising rates of obesity ^3^.

Insulin resistance (IR) is a fundamental pathophysiological feature and a key driver in the development and progression of type 2 diabetes mellitus (T2DM). It is defined as a state of reduced responsiveness of insulin-targeting tissues to physiological levels of insulin, disrupting glucose homeostasis in key metabolic organs such as the liver, skeletal muscle, and adipose tissue ^4^. This impairment compromises glucose uptake, leads to unsuppressed hepatic glucose production, and dysregulates lipid metabolism. In response, pancreatic β-cells augment insulin secretion to compensate for diminished sensitivity, resulting in chronic hyperinsulinemia ^5^. A critical aspect of this condition is selective IR, in which insulin fails to suppress gluconeogenesis but continues to stimulate lipogenesis, thereby exacerbating hyperglycemia, dyslipidemia, and hepatic steatosis ^4,6^. Over time, the compensatory hyperinsulinemia and the glucotoxic environment contribute to β-cell exhaustion and dysfunction, culminating in persistent hyperglycemia and the onset of overt T2DM ^5^.

The molecular mechanisms underpinning IR are complex and multifactorial. Ectopic lipid accumulation in liver and muscle is a central player, as lipid intermediates like diacylglycerol (DAG) can activate protein kinase C (PKC) isoforms, leading to the inhibition of insulin receptor signaling ^4,7^. This mechanism is strongly supported by human studies, in which hepatic DAG content has been identified as the best predictor of hepatic IR ^8^. Other prominent theories involve chronic low-grade inflammation, endoplasmic reticulum (ER) stress, mitochondrial dysfunction, and hexosamine biosynthesis pathway flux, all of which can converge to impair the insulin signaling cascade ^4,5,9^. Understanding the intricate interplay between IR and subsequent β-cell failure is crucial for devising effective therapeutic strategies. While current treatments often focus on glycemic control, emerging approaches aim to directly restore insulin sensitivity and preserve β-cell function, underscoring the pivotal role of overcoming IR in the management and potential reversal of T2DM^4,5,10^.

Emerging evidence underscores the critical role of the gut microbiota in the development of diet-induced IR. Studies have shown that germ-free mice fed a high-fat diet (HFD) exhibited less weight gain and did not develop IR compared to conventional mice fed the same diet ^11^. This suggests that diet-induced obesity may promote IR through mechanisms mediated by the gut microbiota. Specifically, the gut microbiota ferments undigested dietary components to produce short-chain fatty acids (SCFAs), which can enhance monosaccharide absorption in the intestine, thereby increasing hepatic lipogenesis and contributing to IR ^12^. In addition, gut microbes modulate the expression of sodium-glucose cotransporter 1 (SGLT-1) in the small intestine and promote angiogenesis in the intestinal villi, thereby enhancing glucose uptake and energy harvest ^12,13^. The microbiota also influences host appetite and satiety by regulating the secretion of gut hormones such as peptide YY (PYY) and glucagon-like peptide-1 (GLP-1), and can directly interact with the central nervous system to affect feeding behavior ^13,14^.

Furthermore, HFD-induced dysbiosis can compromise intestinal barrier integrity, facilitating the translocation of microbial products ^15^. A key example is the enrichment of gut bacteria that produce lipopolysaccharide (LPS), a significant inflammatory factor ^16^. LPS binds to receptors such as CD14 on monocytes and macrophages, activating immune cells and triggering the release of pro-inflammatory cytokines such as IL-6 and TNF-α. This inflammatory response is implicated in the pathogenesis of obesity, IR, and type 2 diabetes ^17^.

Given this pathophysiology, strategies to modulate the gut microbiota present promising therapeutic avenues. Probiotics have demonstrated beneficial effects on glucose metabolism by restoring microbial balance, enhancing SCFA production, suppressing inflammation, and improving gut barrier integrity ^18^. A systematic review and meta-analysis of 22 randomized controlled trials (involving 2,218 participants) confirmed that probiotics can reduce HbA1c, fasting blood glucose, and HOMA-IR while improving insulin sensitivity without altering body mass index^19^. Another established therapeutic class for T2DM is α-glucosidase inhibitors (AGIs), such as acarbose, miglitol, and voglibose ^20^. These drugs lower postprandial blood glucose by competitively inhibiting carbohydrate-hydrolyzing enzymes in the small intestine ^21^.

Given the established role of AGIs in managing postprandial glucose and the emerging evidence for probiotic benefits in metabolic health, we sought to investigate the efficacy of a probiotic strain with AGI activity. The strain *L. plantarum* VB165 was selected from a panel of isolates due to its potent α-glucosidase inhibitory activity *in vitro* (data not shown). We hypothesized that *L. plantarum* VB165, by virtue of its intrinsic α-glucosidase inhibitory activity and potential probiotic properties, would ameliorate high-fat diet-induced IR and associated metabolic disorders by modulating gut microbiota, strengthening the intestinal barrier, and reducing systemic inflammation.

## Materials and Methods

### Bacterial strain and culture conditions

The bacteria strain *Lactiplantibacillus plantarum* VB165 was isolated from stool sample of a health volunteer without underlying metabolic diseases. The strain was cultured in de Man, Rogosa and Sharpe Medium (MRS, Hopebio, Qingdao, China) agar anaerobically overnight. For fermentation, the strain was inoculated into 100 mL MRS medium or Simulated small intestinal fluid (Coolaber, Beijing, China) and incubated anaerobically at 70 rpm for 24 h. For oral gavage, bacterial suspensions were diluted in PBS (Solarbio, Beijing, China) to a final concentration of 10^9^ CFU/0.2 mL.

### Test for α-Glucosidase Inhibitory Activity

The fermentation culture (1 mL) was centrifuged at 10,000 rpm to obtain the supernatant. α-glucosidase activity assay was performed in a 96-well plate with a total reaction volume of 100 μL. First, 25 μL of the substrate 4-Nitrophenyl α-D-glucopyranoside (PNPG, 10 mmol/L, Solarbio) and 25 μL of the sample were mixed and incubated at 37°C for 10 min. Then, 50 μL of α-glucosidase (40 U/L, Sigma-Aldrich, St. Louis, USA) was added, and the mixture was incubated at 37°C for 30 min. The reaction was stopped by adding 100 μL of Na₂ CO₃ (0.1 mol/L).

Absorbance at 405 nm was measured using a microplate reader (Biotek, Winooski, Vermont, USA). The absorbance of the sample was calibrated using a sample blank control (in which 50 μL of PBS replaced the α-glucosidase). For the negative control (no inhibitory sample), 25 μL of PBS was used to replace the sample. For the negative blank control (no enzyme activity), 25 μL of PBS replaced the sample and 50 μL of PBS replaced the α-glucosidase. The inhibition rate was calculated, and strains exhibiting an inhibition rate > 40% were selected for further testing.

The α-glucosidase inhibition rate was calculated using the following formula: α-Glucosidase Inhibition Rate (%) = (1 - (A - B) / (C - D)) × 100%

Whereas:

A is the absorbance of the sample.

B is the absorbance of the sample blank control.

C is the absorbance of the negative control.

D is the absorbance of the negative blank control.

### Animals, Feeding Procedures, and study design

Male C57BL/6 mice (Animal Center, Hangzhou Normal University) aged 6∼8 weeks were randomly divided into 4 groups (n=10 per group, housed 5 per cage).

Normal control group (NCD): fed a standard chow diet and water.

VB165 group (NCD+VB165): fed a standard chow diet and supplementation with *L. plantarum* VB165 (10^9^ CFU/day) by oral gavage.

High-Fat Diet group (HFD): fed a High-Fat Diet (60% fat, ReseachDiet, New Brunswick, USA) and water.

High-Fat Diet and VB165 group (HFD+VB165): fed a High-Fat Diet (60% fat) and supplementation with *L. plantarum* VB165 (10^9^ CFU/day) by oral gavage.

Mice were maintained on a 12-h light/dark cycles at 20 ± 2℃. Body weight and food intake were recorded weekly for 16 weeks.

After 16 weeks, mice were fasted for 12 h, weighed, and anesthetized. Blood was collected via orbital puncture, followed by cervical dislocation. Tissues (liver, epididymal fat, and intestine) were rapidly excised, flash-frozen in liquid nitrogen, transported on dry ice, and stored at -80℃. Serum was isolated by centrifuging blood at 3,000 rpm for 10 min.

This study was approved by the Ethics Committee of Hangzhou Normal University (No. 2023-1070).

### Oral Glucose Tolerance Test (OGTT)

At week 15, after a 12-h fast, mice received an oral gavage of glucose solution (2 g/kg body weight). Blood glucose levels were measured at 0, 15, 30, 60, and 120 min. The area under the curve (AUC) was calculated from blood glucose values over time^22^.

### Insulin Tolerance Test (ITT)

At week 16, after fasting for 2∼5 h, mice were intraperitoneally injected with insulin solution (0.75 IU/kg body weight). Blood glucose levels were measured at 0, 15, 30, 60, and 120 min, and the AUC was calculated ^22^.

### Measurement of Serum Biochemical Parameters

Blood was collected via orbital puncture and placed in Eppendorf tubes. After clotting for 6 hours, serum was separated by centrifugation at 3000 rpm for 10 min. Levels of triglycerides (TG), total cholesterol (TC), high-density lipoprotein cholesterol (HDL-C), and low-density lipoprotein cholesterol (LDL-C) were measured using an automated biochemical analyzer (LABOSPECT 008 α, Hitachi, Tokyo, Japan).

### Detection of Serum Inflammatory Cytokines, LPS, and Insulin

Pre-coated ELISA kits were used to measure inflammatory cytokines (TNF-α, IL-6, IL-10, IL-1β), LPS, and insulin according to the manufacturer’s instructions (Upingbio, Shenzhen, China).

### Homeostatic Model Assessment for Insulin Resistance (HOMA-IR)

HOMA-IR was calculated using the following formula:

HOMA-IR = Fasting blood glucose (mmol/L) × Fasting insulin (μU/mL) / 22.5

### Western Blot

Colon tissues were homogenized in RIPA lysis buffer (KeyGen Biotech, Nanjing, China). Protein concentration was determined using the BCA assay (Thermo Fisher Scientific, Waltham, USA), and samples were adjusted to 30 mg. Proteins were denatured at 95°C for 5 min.

Equal amounts of protein were separated on a 10% SDS-PAGE gel at 120 V for 1 hour. Proteins were transferred to PVDF membranes (EMD Millipore, Billerica, USA) via wet transfer (20% methanol, 1× Tris-glycine buffer) at 180 mA for 1.5 h.

Membranes were blocked with 5% non-fat milk in TBST (Tris-buffered saline with 0.1% Tween-20) for 2 h at room temperature (RT) and then incubated overnight at 4°C with primary antibodies (anti-GAPDH, anti-Occludin, or anti-ZO-1; CST, Danvers, USA). After washing with TBST (3 × 10 min), membranes were incubated with HRP-conjugated secondary antibodies (Proteintech, Wuhan, China) for 1 h at RT.

Signals were visualized using ECL substrate (Thermo Fisher Scientific, Waltham, USA) and quantified with ImageJ 1.47 (https://imagej.nih.gov/ij/). Band intensities were normalized to GAPDH.

### Histological Studies

Tissues were fixed in 10% formaldehyde in PBS (pH 7.0), dehydrated, and embedded in paraffin. Serial paraffin sections (5 μm thick) were prepared for staining and immunohistochemistry ^23^.

#### Hematoxylin and Eosin (H&E) Staining

Tissue sections were baked at 60°C for 25 min, deparaffinized in xylene (three changes, 10, 8, and 8 min), and then sequentially dehydrated in 90%, 80%, and 70% ethanol (3 min each). After washing with ddH₂ O three times (3 min each), sections were stained with hematoxylin, rinsed with water, differentiated with hydrochloric acid-ethanol, and rinsed again. Eosin staining was applied, followed by washing with tap water for 10 min. Sections were dehydrated in absolute ethanol (three changes) and cleared in xylene (three changes). Finally, sections were air-dried and mounted with resin.

#### Oil Red O Staining

Tissue sections were washed with ddH₂ O to remove embedding medium, rinsed with 60% isopropanol for 2 min, and stained with Oil Red O (Sigma-Aldrich) working solution for 5 min in the dark. Sections were then rinsed with 60% isopropanol for 5∼10 sec for color adjustment, immediately washed with ddH₂ O, counterstained with hematoxylin for 1 min, and rinsed again with ddH₂ O for color development. Sections were mounted with glycerol gelatin and observed under a microscope for imaging.

#### Immunohistochemistry

Slides were deparaffinized, rehydrated, and washed with PBS. Sections were blocked with goat serum albumin for 10 min and then incubated overnight at 4°C with mouse anti-Occludin or anti-ZO-1. The next day, after washing with PBST, sections were incubated with biotinylated anti-mouse antibody (1:5000) for 10 min at RT, followed by HRP-streptavidin (1:500) for 10 min. After washing, samples were counterstained with DAB solution (Sigma-Aldrich).

### Gut Microbiota Analysis

DNA was extracted from stool samples using the DNeasy DNA Extraction Kit (Qiagen, Duesseldorf, Germany). The 16S V3∼V4 region was amplified with primers 338F (5’-ACT CCT ACG GGA GGC AGC A-3’) and 806R (5’-GGA CTA CHV GGG TWT CTA AT-3’) ^24^. Library preparation and sequencing were performed using the PE250 protocol on the Illumina NovaSeq platform (Illumina, San Diego, USA).

Raw sequencing reads were quality-controlled using Fastp and analyzed with Qiime2 (Greg Caporaso Lab, Flagstaff, USA). Paired-end reads were merged, quality-filtered, dereplicated, and clustered into operational taxonomic units (OTUs) at 97% similarity using VSEARCH ^25^. Taxonomic classification was performed against the SILVA 138.1 database ^26^. Taxonomic composition was visualized using the Qiime taxa barplot tool ^27^. Feature tables and taxonomy annotations were exported in BIOM format and converted to TSV for detailed analysis ^28^.

Species composition and differential abudance analysis were performed using QIIME2 and R 3.6.3, followed by graphical representation ^27^. LEfSe software was employed to conduct linear discriminant analysis (LDA) to assess the impact of species abundance on intergroup microbiota differences, and identifying characteristic microbial taxa for each group. The comparative analysis between groups was performed by STAMP^29^, with the Wilcoxon rank-sum test applied to identify differentially abundant species.

### Statistical Analysis

Statistical analyses were performed using GraphPad Prism 6.01 software (GraphPad Software, La Jolla, CA, USA). One-way ANOVA was used to determine *p* value. All data were expressed as mean ± standard deviation (SD).

## Results

### L. plantarum VB165 exhibited α-Glucosidase Inhibitor (AGI) activity

The culture supernatants of VB165 inhibited α-glucosidase enzyme activity both under normal culture condition (MRS) and in simulated intestinal fluid (Figure S1). Maltose and sucrose, substrates of human α-glucosidase, were used as the carbon source for VB165. Intraperitoneal administration of the bacterial suspension of VB165 at 5×10^9^ CFU did not cause any significant change in the body weight of the mice throughout the observation period, indicating that VB165 is non-pathogenic (Figure S2). The VB165 strain was sensitive to most commonly used antibiotics such as ampicillin, erythromycin, amoxicillin, clarithromycin, and levofloxacin (Table S1).

**Figure S1.**
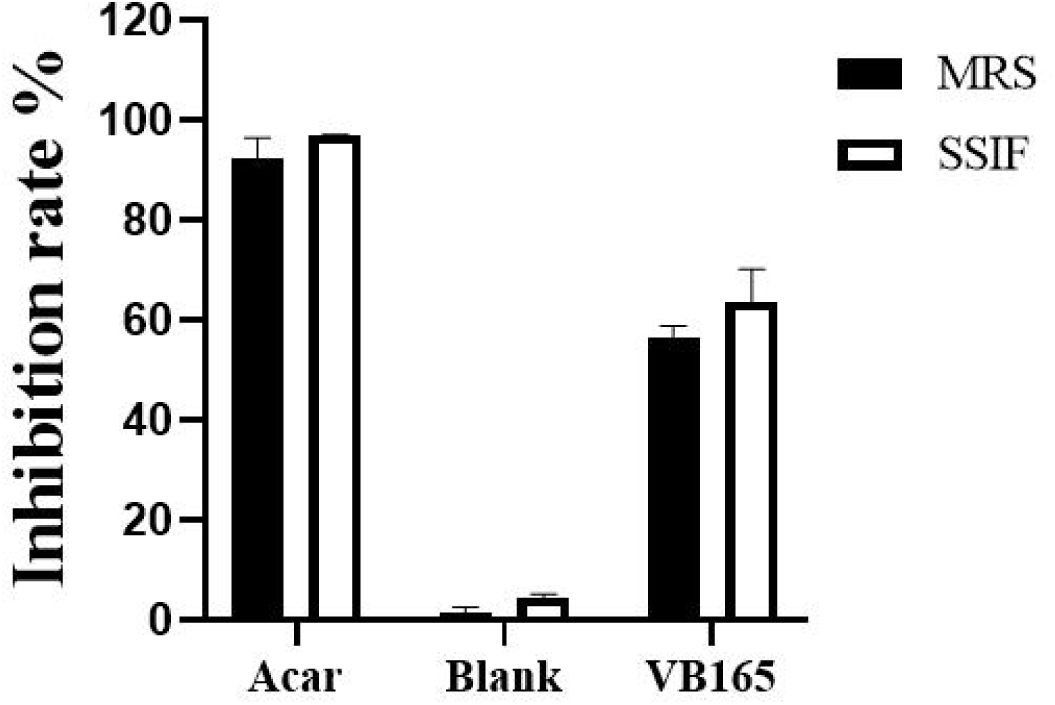
The *L. plantarum* VB165 exhibited α-glucosidase inhibitory activity. Stain VB165 was cultured in MRS and Simulated small intestinal fluid (SSIF). The supernatants of fermentation culture were used to test α-glucosidase inhibitory activities. Acarbose at 1 mg/mL was used as positive control.

**Figure S2.**
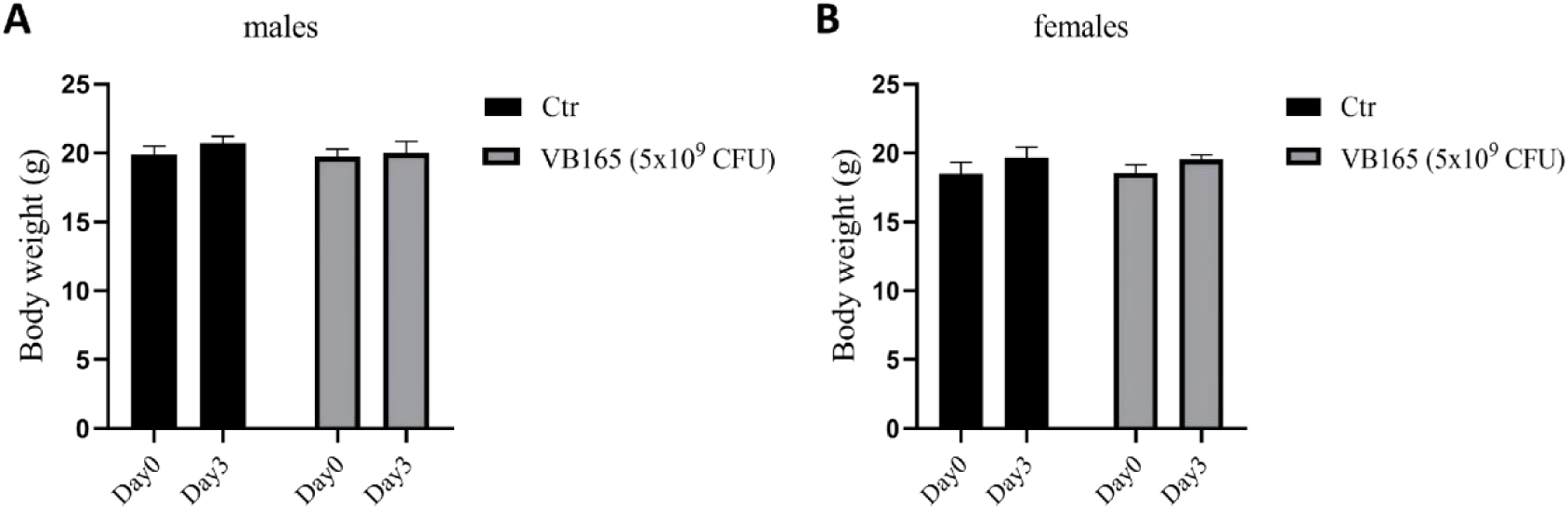
Intraperitoneal challenge with strain VB165 in mice. Male (A) and female (B) ICR mice weighing between 18 g and 22 g were selected. In the control group (N=10, with an equal number of males and females), each mouse received an intraperitoneal injection of 0.3 mL physiological saline. In the experimental group (N=10, with an equal number of males and females), each mouse was intraperitoneally injected with 0.3 mL of fresh bacterial suspension, (5 × 10⁹ CFU). The mice were observed for three days, and changes in behavior as well as fluctuations in body weight were recorded.

**Table S1.** Antimicrobial resistance phenotype of *L. plantarum* VB165 The resistance phenotype of the strain was tested as recommended by the European Food Safety Authority (EFSA). The minimum inhibitory concentration (MIC) for each antimicrobial was evaluated using the microdilution method as described in the international standard ISO 10932:201020.

| Antibiotics | MIC<sup>a</sup> (mg/L) | Cut-off<sup>b</sup> (mg/L) | Interpretation |
| --- | --- | --- | --- |
| Ampicillin | 0.5 | 2 | Sensitive |
| Kanamycin | 128 | 64 | Resistance |
| Erythromycin | 0.5 | 1 | Sensitive |
| Tetracycline | 2 | 32 | Sensitive |
| Chloramphenicol | 2 | 8 | Sensitive |
| Amoxicillin | 0.03 | 2 | Sensitive |
| Clindamycin | 0.25 | 2 | Sensitive |
| Streptomycin | 128 | NR<sup>c</sup> | NR |

<sup>a</sup>Minimum inhibitory concentrations for *L. plantarum* VB165 determined in this study
<sup>b</sup>Microbiological cut-off values for *Lactobacillus plantarum/pentosus* according to EFSA, 2012
<sup>c</sup>NR, not required

### L. plantarum VB165 supplementation attenuates high-fat diet-induced weight gain and adiposity

Feeding mice with HFD led to increased body weight. By week 16, the body weight and total weight gain of HFD mice were significantly higher than those of the NCD group. Supplementation with *L. plantarum* VB165 significantly attenuated weight gain compared to the HFD group (Figure 1A). In contrast, there was no significant difference in energy intake between the HFD and HFD+VB165 groups (Figure 1B), indicating that the difference in weight gain was not due to variations in energy intake.

**Figure 1.**
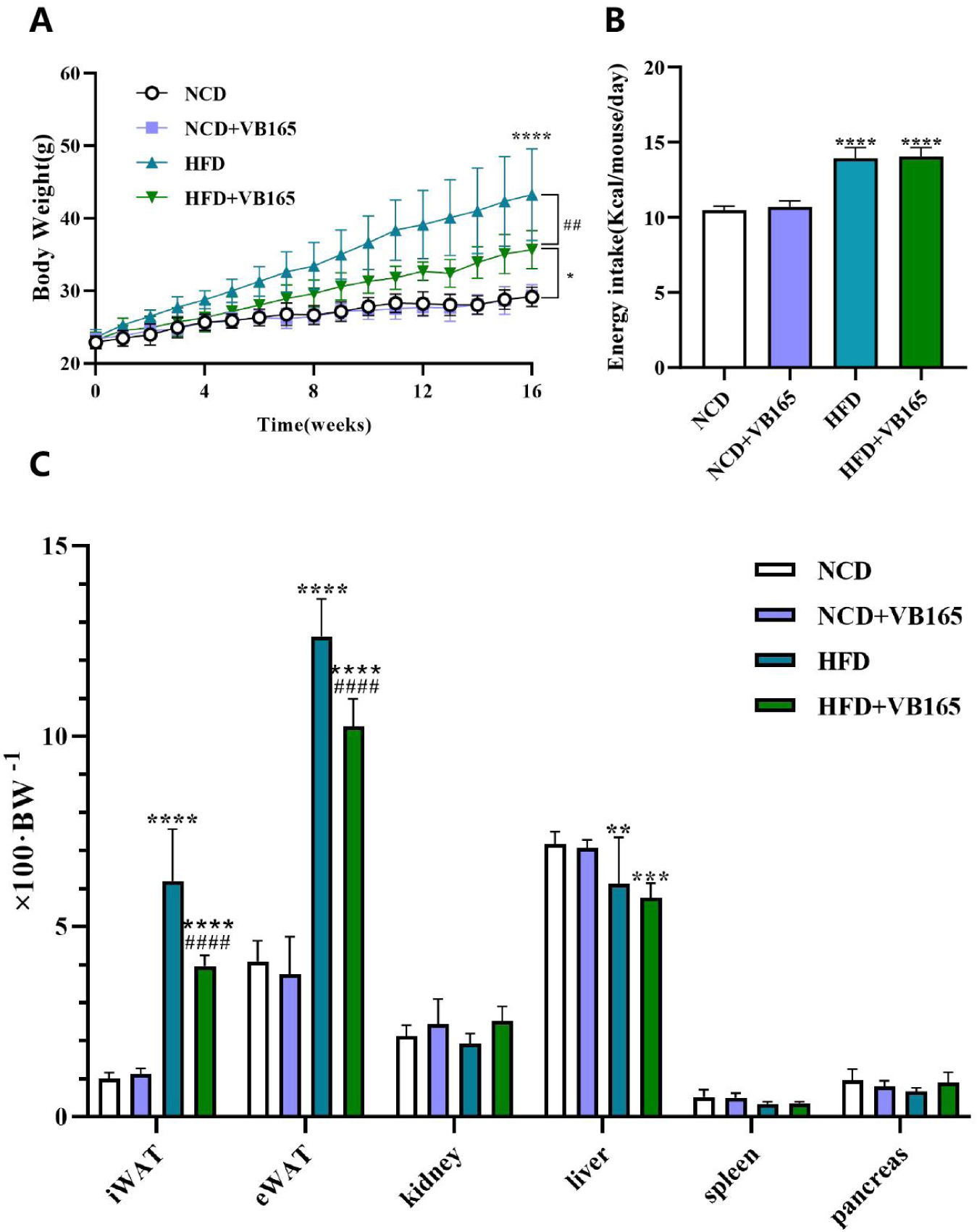
Effects of *L. plantarum* VB165 supplementation on body weight, energy intake and organ index in HFD-fed mice. (A) Body weight curves over 16 weeks; (B) Cumulative energy intake; (C) Organ index (% of body weight). Data were expressed as mean ± SD. One-way ANOVA for multiple comparisons: * p < 0.05; ** p < 0.001; *** p < 0.001; **** p < 0.0001(compared with the NCD group). # p < 0.05; ## p < 0.001; ### p < 0.001; #### p < 0.0001(compared with the HFD group).

Consistent with the body weight changes, HFD significantly increased epididymal white adipose tissue (eWAT) and inguinal white adipose tissue (iWAT) indices compared to NCD groups (Figure 1C), reflecting obesity-related fat deposition. A decreased liver organ index was also observed in HFD groups. Intervention with VB165 significantly decreased eWAT and iWAT indices, suggesting a mitigation of adiposity.

### L. plantarum VB165 supplementation Improves HFD-Induced IR

HFD induced significant increases in fasting blood glucose levels after 8 weeks (Figure S2). At week 16, the HFD group exhibited severe glucose intolerance in the oral glucose tolerance tests (OGTT, Figure 2A). The area under the curve (AUC) for blood glucose was significantly larger in the HFD groups than in the NCD group, indicating impaired glucose tolerance. However, VB165 supplementation significantly reduced postprandial blood glucose levels and the corresponding AUC compared to the HFD group (Figure 2B).

**Figure 2.**
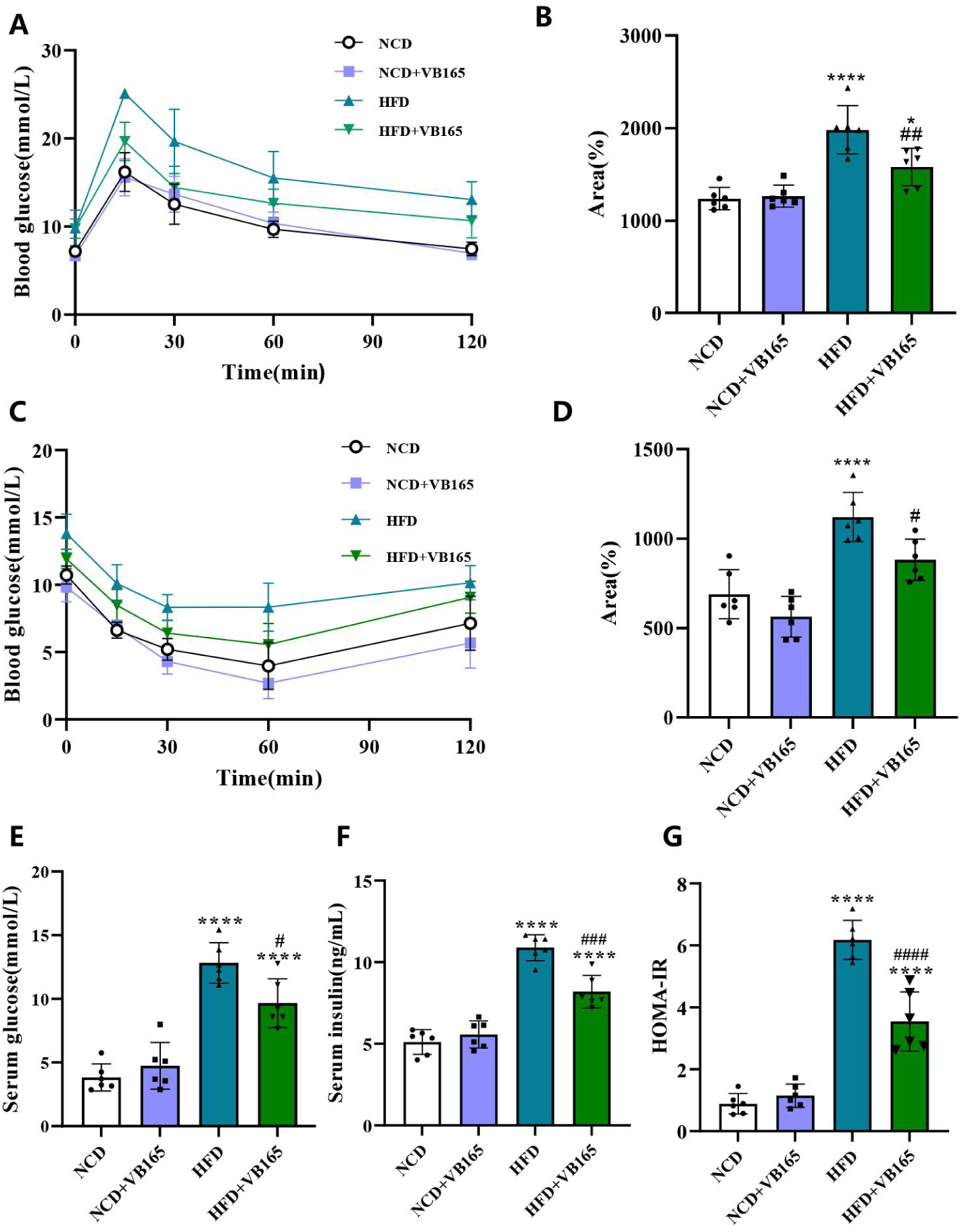
Effects of *L. plantarum* VB165 supplementation on glucose metabolism and IR in HFD-fed mice. (A) OGTT curve; (B) AUC for OGTT; (C) ITT curve; (D) AUC for ITT; (E) Fasting blood glucose levels at week 16; (F) fasting serum insulin levels at week 16; (G) Homeostatic model assessment for insulin resistance (HOMA-IR). One-way ANOVA for multiple comparisons: * p < 0.05; ** p < 0.001; *** p < 0.001; **** p < 0.0001 (compared with the NCD group). # p < 0.05; ## p < 0.001; ### p < 0.001; #### p < 0.0001 (compared with the HFD group).

The results of the intraperitoneal insulin injection test (ITT) were shown in Figure 2C. Mice in the two HFD groups showed higher blood glucose levels at all time points after insulin injection compared to NCD groups (Figure 2C). Probiotic VB165 supplementation in HFD-fed mice significantly reduced the AUC of blood glucose (Figure 2D), suggesting improved insulin sensitivity.

Additionally, compared to the NCD groups, the HFD group exhibited significantly higher fasting blood glucose (Figure 2E), fasting insulin (Figure 2F), and the insulin resistance index (HOMA-IR; Figure 2G), indicating pronounced IR. Importantly, VB165 supplementation significantly reduced these HFD-induced metabolic disturbance, including hyperglycemia, hyperinsulinemia, and elevated HOMA-IR.

### L. plantarum VB165 Improves HFD-Induced dyslipidemia and hepatic steatosis

High-fat feeding significantly increased serum levels of TC, TG, and LDL-C in the HFD group (Figures 3A-C), reaching averages of 4.5 mmol/L, 1.6 mmol/L, and 0.5 mmol/L, respectively. No significant differences were observed in HDL-C levels among the four groups (Figure 3D). Compared to the HFD group, the HFD+VB165 group showed significantly lower serum TC, TG, and LDL-C levels, at 3.9 mmol/L, 1.2 mmol/L, and 0.35 mmol/L, respectively.

**Figure 3.**
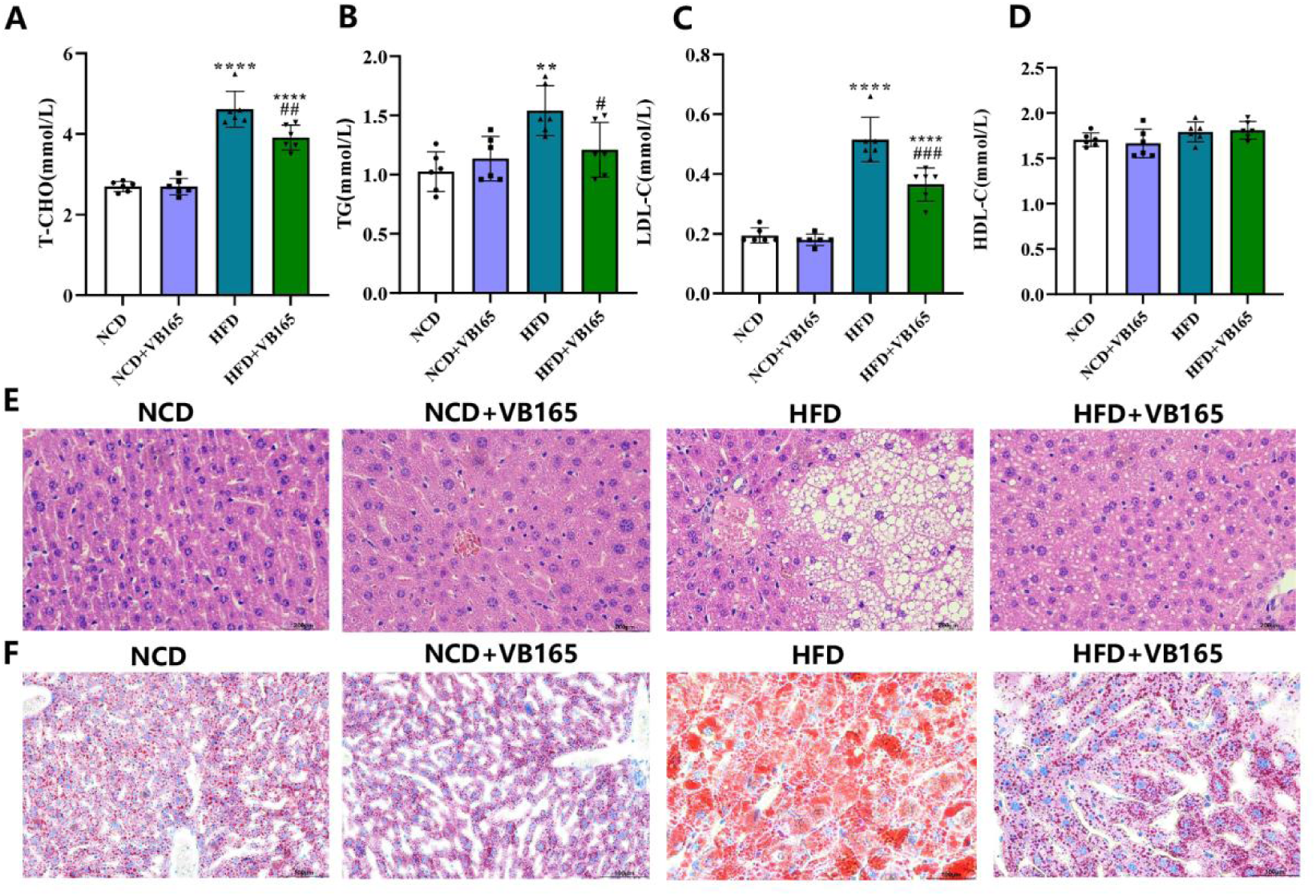
Effects of *L. plantarum* VB165 supplementation on serum lipids and hepatic steatosis in HFD-fed mice. Serum levels of (A) T-CHO, (B) TG, (C)LDL-C, and (D) HDL-C at week 16; (E) Representative liver sections stained with H&E; (F) Representative liver sections stained with Oil Red O. One-way ANOVA for multiple comparisons: * p < 0.05; ** p < 0.001; *** p < 0.001; **** p < 0.0001(compared with the NCD group). # p < 0.05; ## p < 0.001; ### p < 0.001; #### p < 0.0001(compared with the HFD group).

Histological analysis of liver tissue sections revealed that, compared to the NCD group, the HFD group exhibited numerous prominent lipid vacuoles (indicative of lipid droplets) in hepatocytes, while fewer lipid droplets were observed in the HFD+VB165 group (Figure 3E). Similarly, HFD-fed mice had significantly larger epididymal adipocytes than the NCD group. The average adipocyte area was approximately 44,000 μm² in the HFD group and 21,600 μm² in the HFD+VB165 group, indicating a significant reduction in adipocyte size with VB165 intervention.

### L.plantarum VB165 enhances intestinal barrier function and reduces systemic inflammation

High-fat feeding is known to increase intestinal permeability. Therefore, we used immunohistochemistry and Western blotting to detect the expression of tight junction proteins ZO-1 and Occludin in the colon. The results (Figures 4A, B) showed that ZO-1 and Occludin expression was significantly reduced in the HFD group, while the HFD+VB165 group exhibited significantly higher levels of these proteins compared to the HFD group.

**Figure 4.**
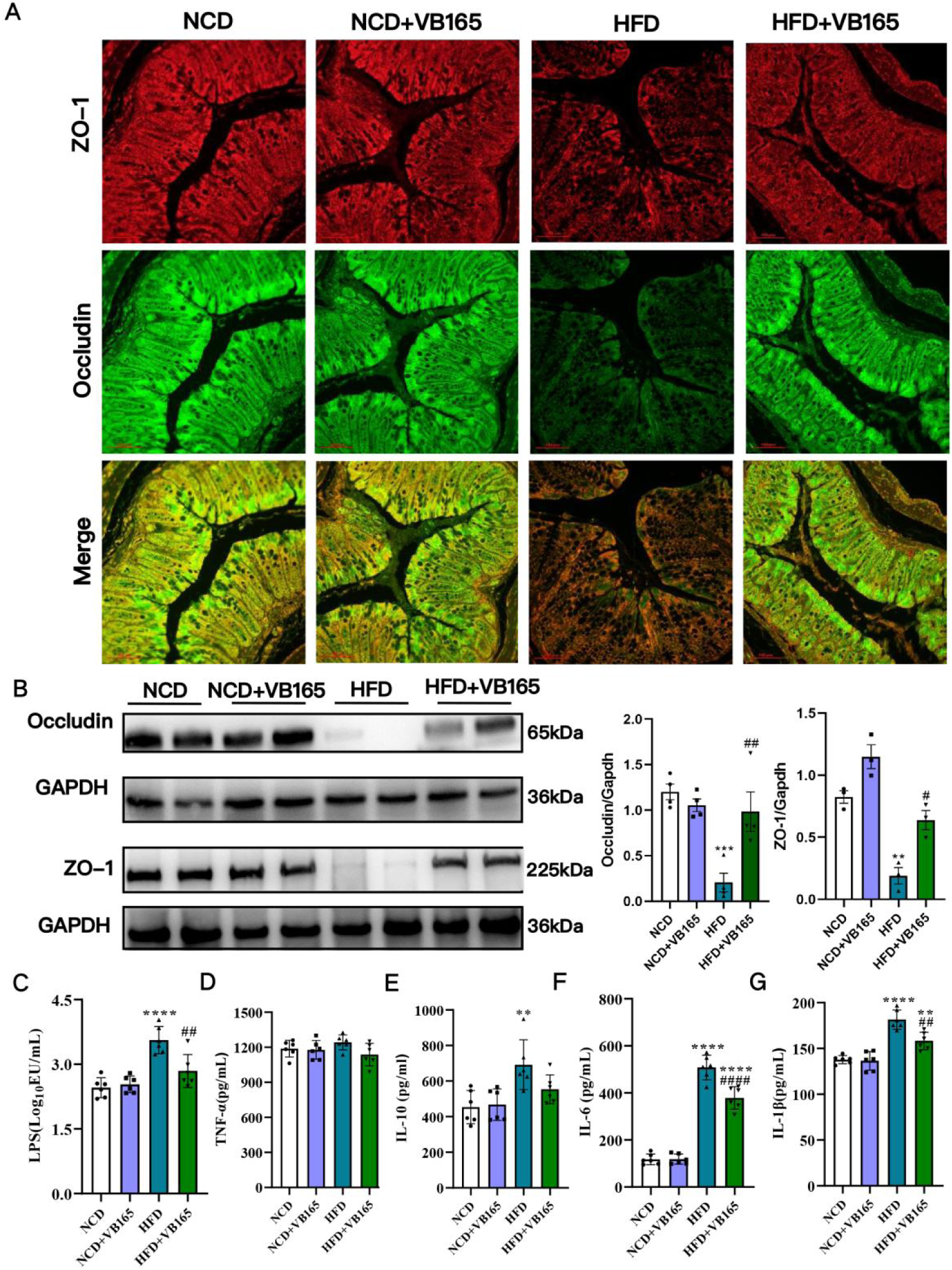
Effects of *L. plantarum* VB165 supplementation on intestinal tight junction proteins and systemic inflammation in HFD-fed mice. (A) Immunohistochemical staining of ZO-1 and Occludin in colon tissue; (B) Western blot analysis of ZO-1 and Occludin protein expression. Serum levels of (C) LPS, (D) TNF-α, (E) IL-10, (F) IL-6, and (G) IL-1β at week 16. One-way ANOVA for multiple comparisons: * p < 0.05; ** p < 0.001; *** p < 0.001; **** p < 0.0001(compared with the NCD group). # p < 0.05; ## p < 0.001; ### p < 0.001; #### p < 0.0001(compared with the HFD group).

Since LPS, a cell wall component released by gram-negative gut bacteria, can activate TLR4 signaling and induce inflammation, we measured its serum levels. As shown in Figure 4C, HFD increased serum LPS levels, but this increase was reduced in the HFD+VB165 group. No significant differences were observed in serum TNF-α levels among the four groups (Figure 4D). However, HFD induced elevated levels of IL-6, IL-10, and IL-1β (Figure 4E-G). Notably, VB165 supplementation significantly reduced levels of IL-6 and IL-1β but not IL-10. This indicates that *L. plantarum* VB165 intervention can partially alleviate HFD-induced low-grade systemic inflammation.

### L plantarum VB165 supplementation modifies gut microbiota composition in HiFD Mice

High-fat feeding did not affect the α-diversity of gut microbiota in mice, while VB165 supplementation significantly increased the Chao1 index, Simpson index, and the number of Observed species (Figure 5A-D). Principal component analysis (PCA) was used to assess β-diversity (Figure 5E). The results showed that HFD significantly alters gut microbiota composition, forming a separate cluster from the NCD groups. VB165 treatment modified, but did not fully reverse, these HFD-induced shifts.

**Figure 5.**
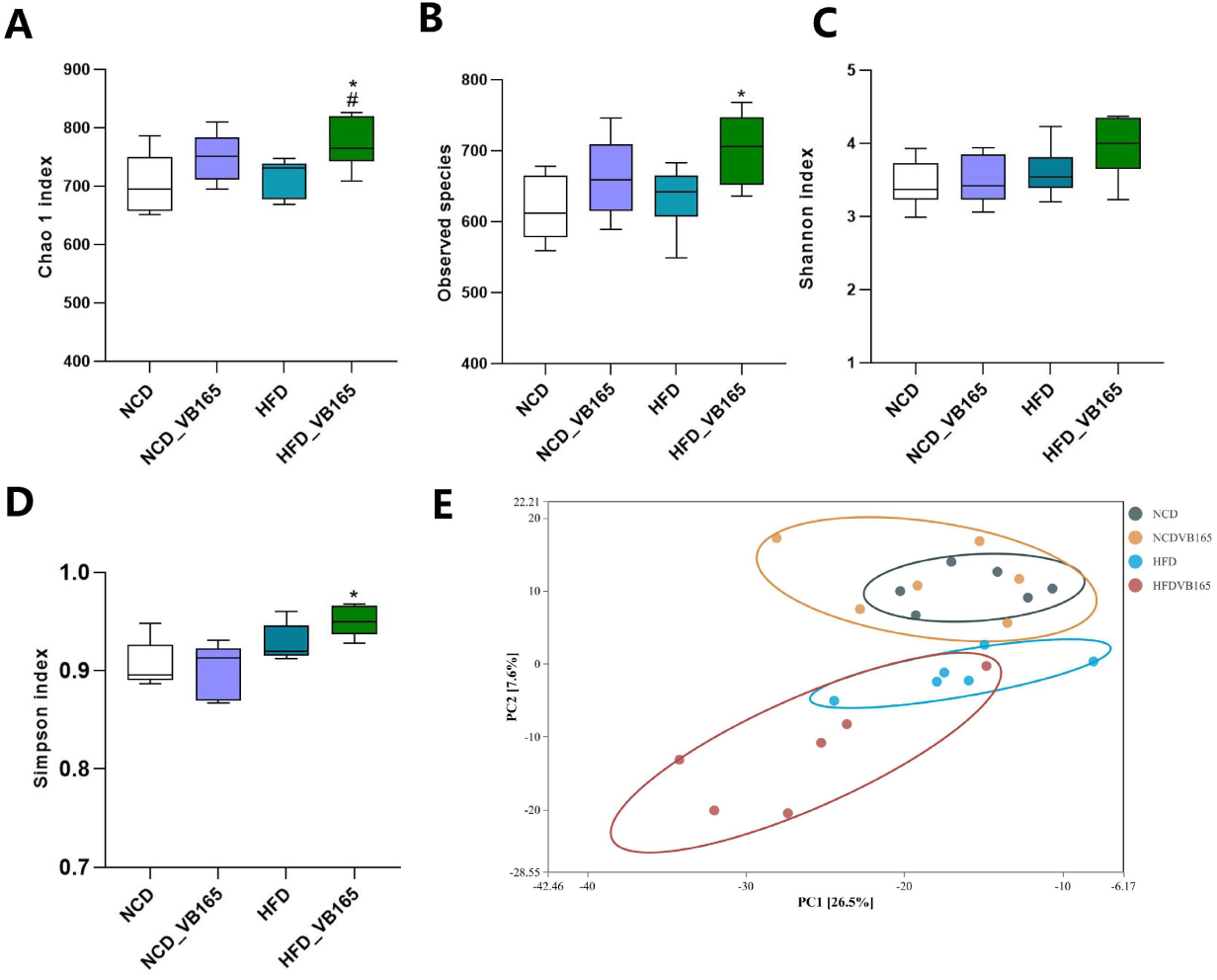
Effects of *L. plantarum* VB165 supplementation on gut microbiota in HFD-fed mice. α-diversity indices including (A) Chao 1, (C) Shannon, and (D) Simpson index, along with (B) observed species were compared. (E) Principal component analysis (PCA) was used to assess β-diversity. One-way ANOVA for multiple comparisons: * p < 0.05; ** p < 0.001; *** p < 0.001; **** p < 0.0001(compared with the NCD group). # p < 0.05; ## p < 0.001; ### p < 0.001; #### p < 0.0001(compared with the HFD group).

At the genus level (Figure 6A), *L. plantarum* VB165 supplementation significantly reduced the abundance of *Coriobacteriaceae*_UCG_002 (a genus of *Coriobacteriaceae*) and *Ileibacterium* in HFD mice, while significantly increasing the abundance of *Faecalibaculum, Lachnospiraceae*_UCG_002, and *Oscillibacter*. Linear discriminant analysis effect size (LEfSe) revealed 25 genera with significant differences among the four groups: 9 enriched in the NCD groups and 16 in the HFD groups. Further analysis (Figures 6B) showed that the NCD group was enriched with *g_Lactobacillus* and *g_Alistipes*. The HFD group was enriched with *Ileibacterium* and *Coriobacteriaceae*_UCG_002, while VB165 supplementation reduced the abundance of these genera. The HFD+VB165 group was enriched with *g_Coprococcus, g_Collidextribacter, g_UCG-009,* and *g_Monoglobus*.

**Figure 6.**
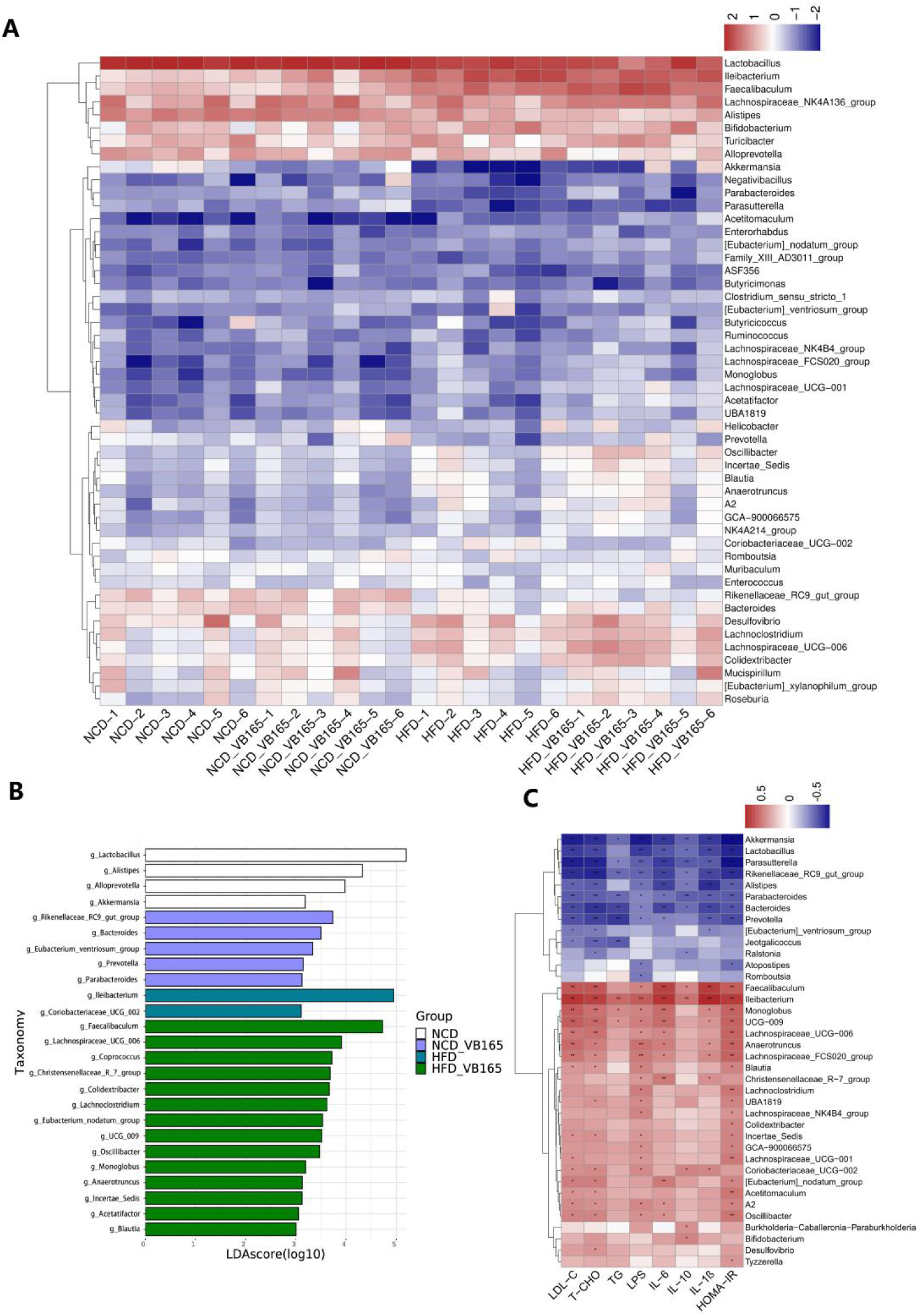
Gut microbiota composition and correlations with metabolic parameters. (A) Heatmap of relative abundance of bacteria genera. (B) LEfSe plot showing 25 taxa that significantly differ among groups (p < 0.05). (C) Spearman correlation heatmap between dominant bacterial genera and metabolic/inflammatory markers.

To investigate the relationships between gut microbiota composition and indicators of metabolic disorders, we performed Spearman correlation analysis using key metrics: lipid metabolism (LDL-C, TC, TG), glucose metabolism (HOMA-IR), and inflammation (LPS, IL-6, IL-10, IL-1β, TNF-α). The results were shown in Figure 6C. The abundance of *Alistipes*, enriched in the NCD group, was negatively correlated with all key metabolic and inflammatory metrics. In contrast, *Ileibacterium* and *Coriobacteriaceae_UCG_002*, enriched in the HFD group, were positively correlated with lipid metabolism metrics (LDL-C and TC) and inflammation-related indicators (LPS and IL-1β). Additionally, *g_Coprococcus*, *g_Collidextribacter, g_UCG-009, and g_Monoglobus*, enriched in the HFD+VB165 group, were negatively correlated with lipid metabolism and inflammation-related metrics.

## Discussion

The escalating global burden of IR and T2DM necessitates novel therapeutic strategies. This study introduces *L. plantarum* VB165 as a unique multi-functional therapeutic candidate, distinguished by its dual-action mechanism that combines direct enzymatic inhibition—potent intrinsic α-glucosidase inhibitor (AGI) activity—with indirect, broad-spectrum probiotic effects including gut microbiota remodeling, intestinal barrier fortification, and systemic anti-inflammatory action. This synergistic combination underpins its potent efficacy against HFD-induced metabolic syndrome.

VB165 exhibited direct AGI activity (Figure S1), a property not universally shared by probiotics. This property likely underlies the significant attenuation of postprandial hyperglycemia observed in the OGTT, as it might competitively inhibit carbohydrate-digesting enzymes in the small intestine, thereby flattening postprandial glucose peaks and reducing the glycemic burden on pancreatic b-cells. By mitigating postprandial glucose excursions, VB165 may alleviate glucotoxicity, a key driver of b-cell dysfunction and peripheral IR, thereby breaking a vicious cycle that exacerbates metabolic dysregulation.

Alongside the AGI activity, VB165 induced systemic metabolic improvements, including attenuated body weight gain and reduced adiposity independent of caloric intake (Figure 1). This aligns with the established role of gut microbiota in regulating host energy harvest and storage ^12^. Notably, the reduction in epididymal and inguinal adipose tissue indices, coupled with decreased adipocyte hypertrophy, suggest that VB165 may promote a metabolically healthier adipose phenotype, potentially enhancing adipokine secretion and lipid buffering capacity. Furthermore, VB165 ameliorated HFD-induced dyslipidemia and ectopic fat accumulation (Figure 3). The reduction in hepatic steatosis is particularly critical, as hepatic lipid intermediates like diacylglycerol (DAG) are key drivers of hepatic IR in humans ^8^.

A pivotal pathway through which VB165 exerts its systemic effects is by reinforcing the gut barrier. The upregulation of colonic tight junction proteins (ZO-1, Occludin) in the HFD+VB165 group was functionally linked to a reduction in circulating LPS and pro-inflammatory cytokines (IL-6, IL-1β). This demonstrates VB165’s role in mitigating HFD-induced “metabolic endotoxemia,” a well-established process where gut-derived LPS triggers TLR4-mediated inflammation that disrupts insulin signaling^17^.

The modulation of the gut ecosystem was central to this process. VB165 enhanced microbial α-diversity and significantly increased community richness (Chao1) and evenness (Simpson index), suggesting a restoration of gut microbiota that is affected by HFD. It specifically reversed HFD-driven dysbiosis by suppressing genera enriched by HFD, such as *Coriobacteriaceae*_UCG_002 and *Ileibacterium* (Figure 5)*. Coriobacteriaceae*_UCG_002 was identified as a risk factor for T2DM ^30^ and correlated with adverse serum lipids ^31^, while *Ileibacterium* is associated with inflammation ^32^. Concurrently, it promoted potentially beneficial genera like *Faecalibaculum* and *Oscillibacter* ^33,34^. While these correlations do not establish causality, they strongly suggest that the metabolic improvements conferred by VB165 are mediated, in part, through these specific microbial shifts. The negative correlations between VB165-enriched taxa (e.g., g_*Coprococcus*) and key metabolic disease markers further support their potential beneficial role.

Importantly, the effects reported here are likely strain-specific and may not be generalizable to other *L. plantarum* strains, highlighting the unique properties of VB165. This underscores the importance of screening and characterizing individual probiotic strains for targeted metabolic applications. To formally establish the causal role of the microbiota in the observed phenotypes, future studies employing fecal microbiota transplantation (FMT) from VB165-treated mice to germ-free or antibiotic-treated recipients would be required. Furthermore, while this study focused on systemic and microbial outcomes, investigating the direct impact of VB165 on hepatic and adipose tissue insulin signaling pathways (e.g., IRS-1/PI3K/Akt) would provide deeper mechanistic insight into how gut-derived signals translate to improved peripheral insulin sensitivity.

In conclusion, *L. plantarum* VB165, a probiotic with inherent α-glucosidase inhibitory activity, significantly alleviates diet-induced metabolic syndrome by concurrently modulating host digestion and gut microbiota ecology. This multi-targeted approach, targeting both a key step in carbohydrate metabolism and the underlying gut-systemic axis of inflammation and dysbiosis, makes it a promising candidate for further development as a nutritional or therapeutic intervention for IR and T2DM.

## Acknowledgement

This work was supported by the Huadong Medicine Joint Funds of the Zhejiang Provincial Natural Science Foundation under Grant No. LHDMZ24H04001.

## Author contributions

J.Yang, D.Q. Chen designed the research; T.Xu, K.X.Jiang, and W.Zhang performed the experiments; Y.C. Yang, Z.M.Du, H.M.Zhou, Y.Hui and S.F.Han contributed to reagents/materials; T.Duan, X.Q Wu and Z.R.Zheng analyzed the data; T.Xu and W.Zhang wrote the initial draft; J.Yang and D.Q. Chen revised and edited the manuscript.

## Conflict of Interests

W. Zhang, X. Wu, Z. Zheng, and Y. Yang are employees of Hangzhou VicrobX Biotech Inc., Ltd., which is the patent holder of the *Lactiplantibacillus plantarum* VB165 strain.

## Reference

1. International Diabetes Federation, I. IDF Diabetes Atlas. In IDF Diabetes Atlas. International Diabetes Federation vol. 11th editi (2025).

2. Jia, W., Chan, J. C. N., Wong, T. Y. & Fisher, E. B. Diabetes in China: epidemiology, pathophysiology and multi-omics. Nat. Metab. 7, 16–34 (2025).

3. Zhou, Y. C., Liu, J. M., Zhao, Z. P., Zhou, M. G. & Ng, M. The national and provincial prevalence and non-fatal burdens of diabetes in China from 2005 to 2023 with projections of prevalence to 2050. Mil. Med. Res. 12, 1–15 (2025).

4. Lee, S. H., Park, S. Y. & Choi, C. S. Insulin Resistance: From Mechanisms to Therapeutic Strategies. Diabetes and Metabolism Journal vol. 46 15–37 (2022).

5. Accili, D., Deng, Z. & Liu, Q. Insulin resistance in type 2 diabetes mellitus. Nat. Rev. Endocrinol. 21, 413–426 (2025).

6. Titchenell, P. M. et al. Direct Hepatocyte Insulin Signaling Is Required for Lipogenesis but Is Dispensable for the Suppression of Glucose Production. Cell Metab. (2016) doi:10.1016/j.cmet.2016.04.022.

7. Samuel, V. T. & Shulman, G. I. Mechanisms for insulin resistance: Common threads and missing links. Cell (2012) doi:10.1016/j.cell.2012.02.017.

8. Kumashiro, N. et al. Cellular mechanism of insulin resistance in nonalcoholic fatty liver disease. Proc. Natl. Acad. Sci. U. S. A. (2011) doi:10.1073/pnas.1113359108.

9. Petersen, M. C. & Shulman, G. I. Mechanisms of insulin action and insulin resistance. Physiological Reviews (2018) doi:10.1152/physrev.00063.2017.

10. Gastaldelli, A. et al. Effect of tirzepatide versus insulin degludec on liver fat content and abdominal adipose tissue in people with type 2 diabetes (SURPASS-3 MRI): a substudy of the randomised, open-label, parallel-group, phase 3 SURPASS-3 trial. Lancet Diabetes Endocrinol. (2022) doi:10.1016/S2213-8587(22)00070-5.

11. Liu, R. et al. Gut microbiome and serum metabolome alterations in obesity and after weight-loss intervention. Nat. Med. (2017) doi:10.1038/nm.4358.

12. Canfora, E. E., Jocken, J. W. & Blaak, E. E. Short-chain fatty acids in control of body weight and insulin sensitivity. Nature Reviews Endocrinology (2015) doi:10.1038/nrendo.2015.128.

13. Torres-Fuentes, C., Schellekens, H., Dinan, T. G. & Cryan, J. F. The microbiota–gut–brain axis in obesity. The Lancet Gastroenterology and Hepatology (2017) doi:10.1016/S2468-1253(17)30147-4.

14. Martin, C. R., Osadchiy, V., Kalani, A. & Mayer, E. A. The Brain-Gut-Microbiome Axis. CMGH (2018) doi:10.1016/j.jcmgh.2018.04.003.

15. Schroeder, B. O. & Bäckhed, F. Signals from the gut microbiota to distant organs in physiology and disease. Nature Medicine (2016) doi:10.1038/nm.4185.

16. Caesar, R. et al. Gut-derived lipopolysaccharide augments adipose macrophage accumulation but is not essential for impaired glucose or insulin tolerance in mice. Gut (2012) doi:10.1136/gutjnl-2011-301689.

17. Boutagy, N. E., McMillan, R. P., Frisard, M. I. & Hulver, M. W. Metabolic endotoxemia with obesity: Is it real and is it relevant? Biochimie (2016) doi:10.1016/j.biochi.2015.06.020.

18. Payahoo, L., Ghalichi, F., Fathi, M. & Ehsani, A. A Systematic Review of the Effect of Probiotics, Prebiotics, and Synbiotics on Gut Microbiota in Type 2 Diabetes Mellitus. Endocrinol. Res. Pract. 29, 267–276 (2025).

19. Ayesha, I. E. et al. Probiotics and Their Role in the Management of Type 2 Diabetes Mellitus (Short-Term Versus Long-Term Effect): A Systematic Review and Meta-Analysis. Cureus 15, (2023).

20. Dahlén, A. D. et al. Trends in Antidiabetic Drug Discovery: FDA Approved Drugs, New Drugs in Clinical Trials and Global Sales. Frontiers in Pharmacology (2022) doi:10.3389/fphar.2021.807548.

21. Ceriello, A. et al. Guideline for management of postmeal glucose in diabetes. Diabetes Res. Clin. Pract. (2014) doi:10.1016/j.diabres.2012.08.002.

22. Nagy, C. & Einwallner, E. Study of in vivo glucose metabolism in high-fat diet-fed mice using oral glucose tolerance test (OGTT) and insulin tolerance test (ITT). J. Vis. Exp. 2018, 1–12 (2018).

23. Xu, Y. et al. Sulforaphane Ameliorates Nonalcoholic Fatty Liver Disease Induced by High-Fat and High-Fructose Diet via LPS/TLR4 in the Gut–Liver Axis. Nutrients (2023) doi:10.3390/nu15030743.

24. Dai, T. et al. Nutrient supply controls the linkage between species abundance and ecological interactions in marine bacterial communities. Nat. Commun. (2022) doi:10.1038/s41467-021-27857-6.

25. Rognes, T., Flouri, T., Nichols, B., Quince, C. & Mahé, F. VSEARCH: A versatile open source tool for metagenomics. PeerJ (2016) doi:10.7717/peerj.2584.

26. Quast, C. et al. The SILVA ribosomal RNA gene database project: Improved data processing and web-based tools. Nucleic Acids Res. (2013) doi:10.1093/nar/gks1219.

27. Bolyen, E. et al. Reproducible, interactive, scalable and extensible microbiome data science using QIIME 2. Nature Biotechnology (2019) doi:10.1038/s41587-019-0209-9.

28. McDonald, D. et al. The Biological Observation Matrix (BIOM) format or: How I learned to stop worrying and love the ome-ome. GigaScience (2012) doi:10.1186/2047-217X-1-7.

29. Parks, D. H., Tyson, G. W., Hugenholtz, P. & Beiko, R. G. STAMP: Statistical analysis of taxonomic and functional profiles. Bioinformatics (2014) doi:10.1093/bioinformatics/btu494.

30. Sun, K., Gao, Y., Wu, H. & Huang, X. The causal relationship between gut microbiota and type 2 diabetes: a two-sample Mendelian randomized study. Front. Public Heal. 11, (2023).

31. Han, H. et al. Depletion of Gut Microbiota Inhibits Hepatic Lipid Accumulation in High-Fat Diet-Fed Mice. Int. J. Mol. Sci. 23, (2022).

32. Li, S. et al. Bisphenol S exposure induces intestinal inflammation via altering gut microbiome. Food Chem. Toxicol. (2024) doi:10.1016/j.fct.2024.114830.

33. Zagato, E. et al. Endogenous murine microbiota member Faecalibaculum rodentium and its human homologue protect from intestinal tumour growth. Nat. Microbiol. (2020) doi:10.1038/s41564-019-0649-5.

34. Li, C. et al. Gut microbiome and metabolome profiling in Framingham heart study reveals cholesterol-metabolizing bacteria. Cell 187, 1834–1852.e19 (2024).

